# Loss of p53 tumor suppression function drives invasion and genomic instability in models of murine pancreatic cancer

**DOI:** 10.1101/2021.12.31.472823

**Authors:** Claudia Tonelli, Astrid Deschênes, Melissa A. Yao, Youngkyu Park, David A. Tuveson

## Abstract

Pancreatic ductal adenocarcinoma (PDA) is a deadly disease with few treatment options. There is an urgent need to better understand the molecular mechanisms that drive disease progression, with the ultimate aim of identifying early detection markers and clinically actionable targets. To investigate the transcriptional and morphological changes associated with pancreatic cancer progression, we analyzed the *Kras*^*LSLG12D/+*^; *Trp53*^*LSLR172H/+*^; *Pdx1-Cre* (KPC) mouse model. We have identified an intermediate cellular event during pancreatic carcinogenesis in the KPC mouse model of PDA that is represented by a subpopulation of tumor cells that express Kras^G12D^, p53^R172H^ and one allele of wild-type *Trp53. In vivo*, these cells represent a histological spectrum of pancreatic intraepithelial neoplasia (PanIN) and acinar-to-ductal metaplasia (ADM) and rarely proliferate. Following loss of wild-type p53, these precursor lesions undergo malignant de-differentiation and acquire invasive features. We have established matched organoid cultures of pre-invasive and invasive cells from murine PDA. Expression profiling of the organoids led to the identification of markers of the pre-invasive cancer cells *in vivo* and mechanisms of disease aggressiveness.

## INTRODUCTION

Pancreatic ductal adenocarcinoma (PDA) is a lethal cancer with a 5 year survival rate of only 10% (1). The main factors that contribute to this poor prognosis are late diagnosis and lack of therapeutic response (2). The genetic drivers of PDA are well characterized: the initiating event is an activating mutation of *KRAS* in the exocrine pancreas, which establishes precursor lesions called pancreatic intraepithelial neoplasia, or PanINs. Up to 33% of pancreata in autopsy series mainly from aged patients contain low-grade PanINs, indicating that these are common in the general population and unlikely to progress to infiltrating carcinoma (3). Subsequent inactivating mutations of tumor suppressor genes (e.g. *CDKN2A, TP53, SMAD4*) are required to drive tumor formation, as evidenced by genetically engineered mouse models of PDA and genetic analysis of human tumor samples (4,5).

The transcription factor *TP53* is mutated in 72% of human PDAs (6). Studies using mouse models have demonstrated that Kras activation in combination with point mutation or deletion of one copy of the *Trp53* gene is sufficient to induce invasive and metastatic pancreatic cancer reminiscent of the human disease (7,8). In particular, the progression from PanIN to invasive PDA is associated with mutation of *TP53* followed by loss of heterozygosity (LOH) of the second *TP53* allele (9).

Here, we have confirmed in a well-established mouse model for PDA that oncogenic Kras promotes the formation of PanINs, which then progress into increasingly more dysplastic stages and develop into carcinomas following inactivation of p53. Importantly, we were able to distinguish the histological changes associated with the initial *Trp53* mutation from the changes driven by the subsequent LOH of the wild-type *Trp53* allele. We have identified markers of pre-invasive cancer cells which retained the wild-type copy of *Trp53* and observed the initiation of tumor invasion in tissue sections. We have shown that *Trp53* LOH is the ultimate requisite step in tumor progression for the acquisition of invasive features. Using organoid models, we investigated the biology of PDA precursor cells and uncovered the transcriptional changes linked with *Trp53* LOH.

## RESULTS

### Tumor-adjacent PanINs retain the wild-type *Trp53* allele

To investigate pancreatic cancer progression, we utilized the *Kras*^*LSLG12D/+*^; *Trp53*^*LSLR172H/+*^; *Pdx1-Cre* (KPC) mouse model in which Kras^G12D^ and p53^R172H^ were expressed in the pancreas by virtue of Cre recombinase under the control of Pdx1 promoter (7,10). We showed by PCR-based genotyping that loss of the wild-type *Trp53* allele occurred *in vivo* in cancer cells isolated from KPC tumors, as previously reported (Fig. 1A) (7,11). However, we observed in all tumors a very faint band corresponding to the wild-type copy of *Trp53*, indicating that it was retained in a small fraction of neoplastic cells.

**Fig. 1.**
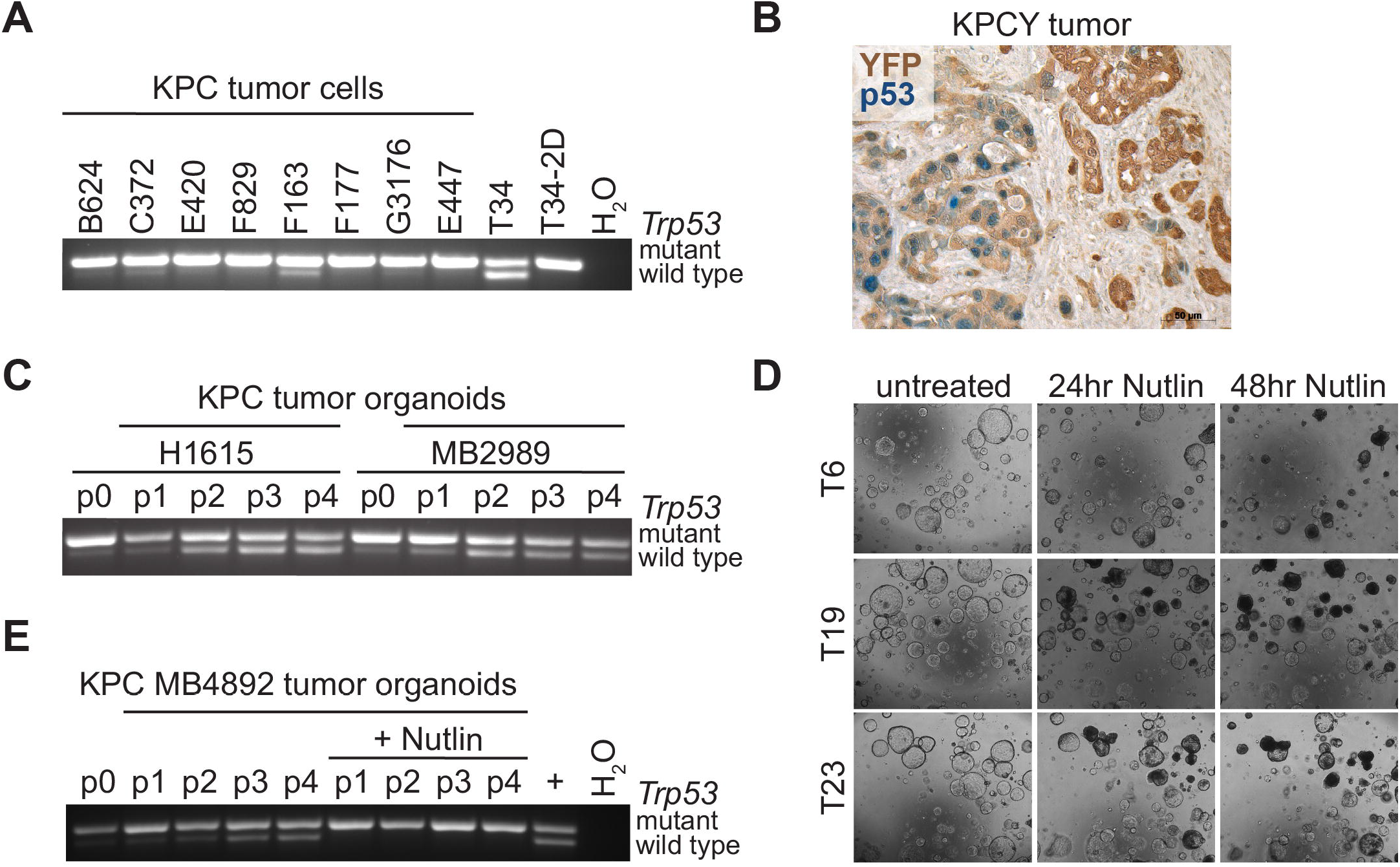
The wild-type *Trp53* allele is lost in most primary KPC tumor cells, but is retained in tumor-adjacent PanINs. **A**. PCR-based genotyping of tumor cells from KPC mice for assessing LOH of the wild-type *Trp53* allele. Tumor ‘T34’ organoids and a 2D line derived from the same T organoids are analyzed as controls. **B**. Immunohistochemical staining for YFP (brown staining) and p53 (blue staining) in a *Kras*^*LSLG12D*^; *Trp53*^*LSLR172H*^; *Pdx1-cre; Rosa26*^*LSLYFP*^ *‘*KPCY’ tumor section. **C**. PCR-based genotyping for assessing the *Trp53* status in tumor cells from KPC mice after isolation (p0) and upon passaging as organoids (p1 to p4). **D**. Pictures of tumor organoids that retain a wild-type *Trp53* allele at 0, 24 and 48 hours after treatment with Nutlin-3a. **E**. PCR-based genotyping for assessing the *Trp53* status in tumor cells from a KPC mouse after isolation (p0) and upon passaging as organoids (p1 to p4) in medium with or without Nutlin-3a.

To corroborate this observation, we analyzed p53 expression in tumor sections of KPCY mice (*Kras*^*LSLG12D/+*^; *Trp53*^*LSLR172H/+*^; *Pdx1-Cre; Rosa26*^*LSLYFP*^), in which all neoplastic pancreatic cells expressed yellow fluorescent protein (YFP) (12). Basal levels of p53 lie below the detection limits of conventional immunohistochemistry (IHC) as wild-type p53 has a rapid turnover until its activity is required to respond to stress signals (7,13,14). Mutations in *Trp53* result in highly stabilized mutant p53 proteins. However, mutation of one *Trp53* allele and retention of the other wild-type *Trp53* allele does not confer increased protein stability probably due to Mdm2-mediated protein degradation (15). *Trp53* LOH is a necessary prerequisite for mutant p53 stabilization and therefore only *Trp53* LOH cells present positive staining for p53 by IHC (7,16). Immunohistochemical staining for YFP and p53 in KPCY tumors showed two populations of YFP-positive neoplastic cells: one with increased nuclear expression of p53 which would represent *Trp53* LOH neoplastic cells, and the other in which p53 was not detected (Fig.1B). Neoplastic cells that did not present p53 accumulation, displayed histological features of PanINs; while cells with detectable p53 expression exhibited features of invasive carcinoma.

Thus, KPC tumors contained neoplastic cells at different stages of tumor progression including PanIN lesions which retained the wild-type *Trp53* allele adjacent to invasive carcinomas.

### Establishment of matched organoid cultures of pre-invasive KPC tumor cells that retain the wild-type *Trp53* allele and invasive KPC tumor cells that have lost the wild-type *Trp53* allele

We previously described the establishment of pancreatic ductal organoid cultures from multiple primary tumors from KPC mice and showed that tumor ‘T’ organoids did not exhibit *Trp53* LOH (17,18). Given that *Trp53* LOH was observed in freshly isolated KPC cancer cells (see Fig. 1A), we sought to determine when *Trp53* LOH cells were lost in organoid culture. We performed PCR-based genotyping of cancer cells after isolation and upon passaging as organoids (Fig. 1C). Tumor cells harboring both a mutant and a wild-type *Trp53* alleles outcompeted the *Trp53* LOH cells in organoid culture in a few passages. The growth advantage was independent of medium or matrix composition and associated with the ploidy status (Suppl. Fig. 1A-C). KPC tumors were enriched in aneuploid and tetraploid cells which were outcompeted in organoid culture by diploid and quasi-diploid cells.

To isolate the *Trp53* LOH cells, we treated the organoids with Nutlin-3a, a small molecule that mediates p53 stabilization by inhibiting the p53-Mdm2 interaction. Nutlin-3a treatment induced the death of the cells that retained a functional wild-type p53 and allowed the isolation of the *Trp53* LOH cells (Fig. 1D, E, Suppl. Fig. 1C, D) (19). Using this strategy, we established matched organoid cultures of KPC cancer cells that retained the wild-type *Trp53* allele (T^m/+^) and KPC cancer cells that had lost the wild-type *Trp53* allele (T^m/LOH^).

### T^m/+^ organoids represent high-grade PanINs

To assess if T^m/+^ organoids could be derived from high-grade PanINs, we compared the transcriptional profiles of T^m/+^ organoids to *Kras*^*G12D/+*^; *Trp53*^*R172H/+*^ (KP) organoids (Table S1). KP and T^m/+^ organoids were both derived from KPC mice and shared the same genotype, however KP organoids were prepared from the pancreata of 2 months old KPC mice that had not yet developed a tumor and presented few low-grade PanIN lesions and areas of ADM, whereas T^m/+^ organoids were derived from the tumor tissue of KPC mice (Suppl. Fig. 2A, B).

Principal component analysis (PCA) depicted the variation among KP and T^m/+^ organoids, which was reflective of mouse variability rather than organoid type (Fig. 2A). Despite the lack of clear transcriptional separation, genes whose levels differed significantly among KP and T^m/+^ organoids were identified. 196 genes were found up-regulated and 160 genes down-regulated in T^m/+^ relative to KP organoids (Fig. 2B, Table S1). Genes that were previously associated with malignant cell transformation controlled cooperatively by loss-of-function p53 and Ras activation were found to be increased in T^m/+^ compared to KP organoids by Gene Set Enrichment Analysis (GSEA) (Fig. 2C) (20). The most notable up-regulated gene was *Cdkn2a*, which encodes for the tumor suppression proteins p16^Ink4a^ and p19^Arf^ through differential exon usage (Fig. 2D) (21). The *Cdkn2a* locus is transcriptionally activated by oncogenic stresses to induce cell cycle arrest or apoptosis: p16^Ink4a^ inhibits Cdk4 activity, while p19^Arf^ mediates the sequestration of Mdm2, resulting in p53 stabilization (22,23). T^m/+^ organoids expressed higher levels of both alternative transcripts. These results indicated that the oncogenic Kras program was more active in T^m/+^ organoids than KP organoids. Furthermore, KP organoids displayed significantly higher expression of normal pancreatic acinar genes such as *Try5, Amy2a1, Amy2a5, Amy2a3, Amy2a2, Amy2a4, Cpb1, Cela2a, Ctrb1* and *Cpa1* (Fig. 2E), suggesting that KP organoids were derived from early PanIN lesions originated from the metaplastic transdifferentiation of acinar to ductal cells. Taken together, our data suggested that KP and T^m/+^ organoids were representative of low-grade and high-grade PanINs, respectively (24,25).

**Fig. 2.**
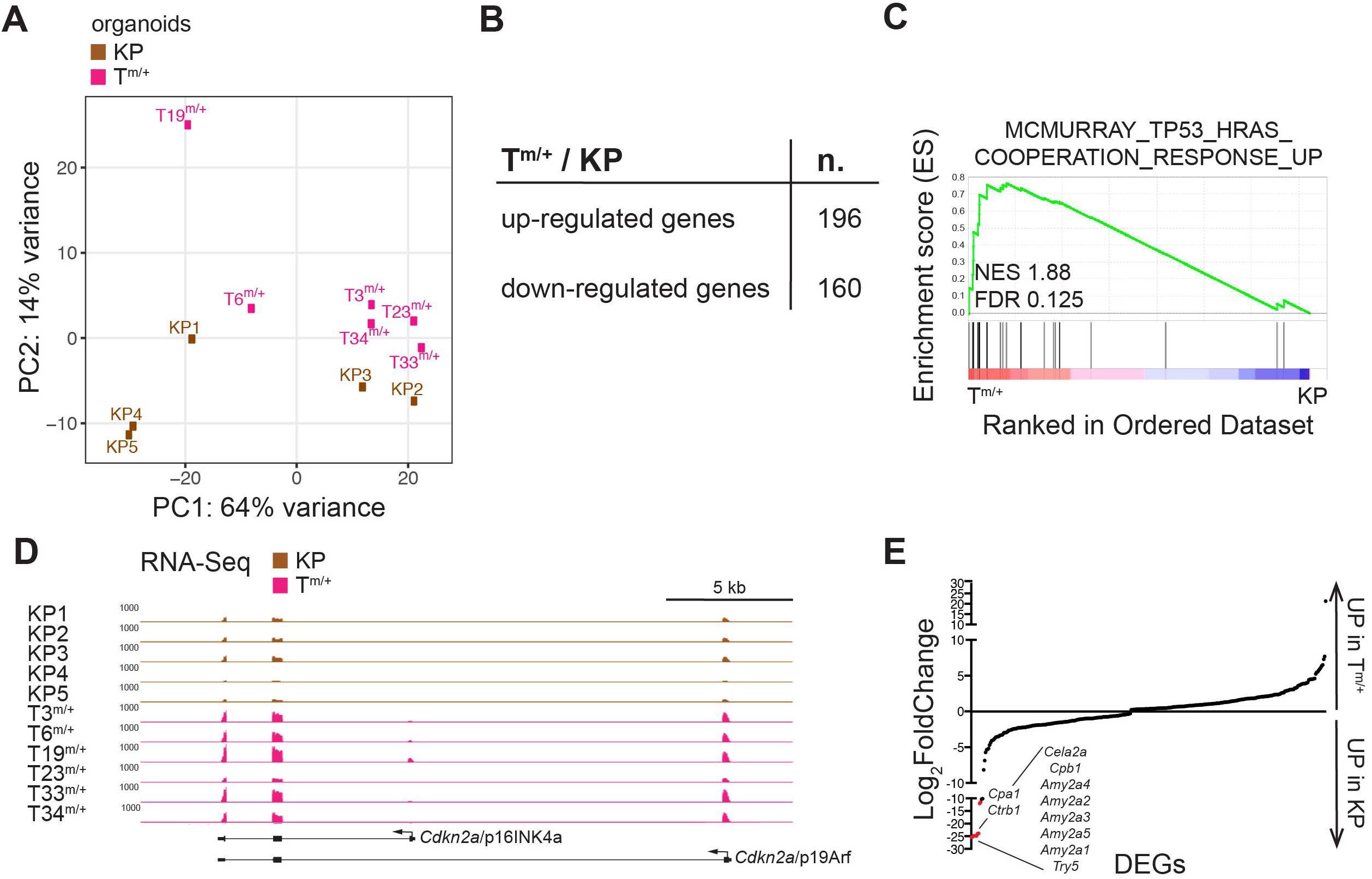
T^m/+^ organoids represent high-grade PanINs. **A**. Principal component analysis of RNA-seq data of KP organoids (n=5) and T^m/+^ organoids (n=6). Samples are colored based on their group: KP organoids in brown and T^m/+^ in pink. **B**. Table of differentially expressed genes between T^m/+^ and KP organoids (up-regulated genes: q-value < 0.05, log2 of fold change > 0; down-regulated genes: q-value < 0.05, log2 of fold change < 0). **C**. GSEA signature ‘MCMURRAY_TP53_HRAS_COOPERATION_RESPONSE_UP’ is induced in T^m/+^ compared to KP organoids. NES, normalized enrichment score; FDR, false discovery rate. **D**. UCSC Genome Browser display of RNA-seq reads in KP and T^m/+^ organoids at the *Cdkn2a* locus. KP organoids tracks are colored in brown and T^m/+^ organoids tracks in pink. **E**. Plot showing the log2 of fold change (T^m/+^ vs KP organoids) for all differentially expressed genes (q-value < 0.05). The top up-regulated genes in KP organoids are indicated.

### *Trp53* LOH is required for PDA development and the acquisition of invasive features

To characterize further the paired organoid models, we established 6 matched organoid cultures of primary KPC tumor cells (Fig. 3A). We found that T^m/+^ organoids expressed higher levels of the wild-type p53 target gene cyclin G1, while T^m/LOH^ organoids acquired stabilization of mutant p53, p19^Arf^ and the DNA damage marker γH2AX in 5 out of 6 lines (26-29). p53 negatively regulates the expression of p19^Arf^; thus elevated p19^Arf^ levels are the result of the interruption of the p53-p19^Arf^ feedback loop (27,28). The accumulation of the DNA damage marker γH2AX is linked to the genome instability that happens as a consequence of p53 inactivation (29).

**Fig. 3.**
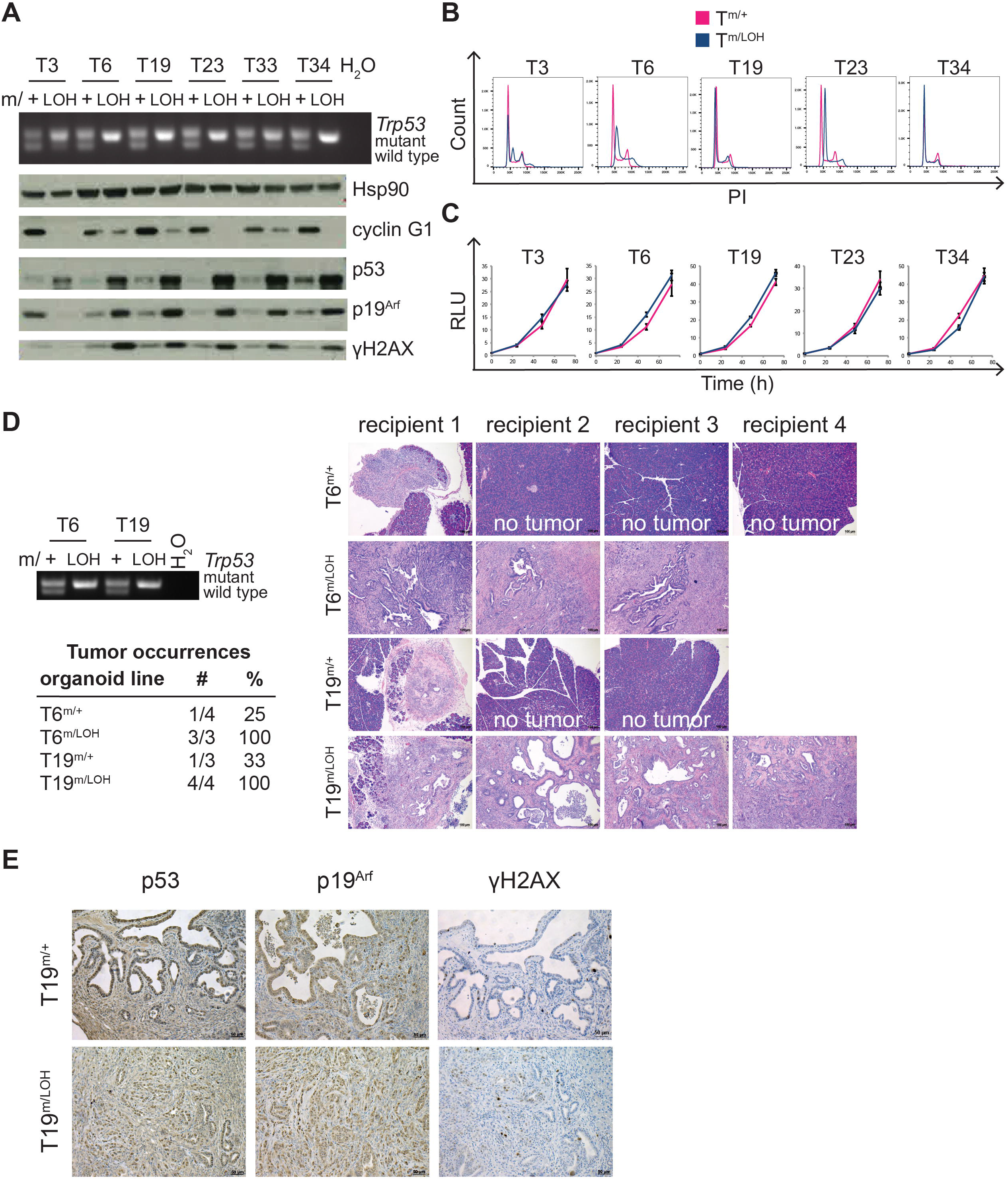
Loss of p53 is the requisite step in tumor progression for the acquisition of invasive features. **A**. PCR-based genotyping of T organoid pairs. T^m/+^ organoids express high levels of the p53 target cyclin G1, while T^m/LOH^ organoids accumulate mutant p53, p19^Arf^ and γH2AX as determined by Western blotting. Loading control, Hsp90. **B**. T^m/+^ organoids (in pink) retain a diploid genome, while some T^m/LOH^ organoids (in blue) show slightly higher DNA content as determined by Propidium Iodide staining. **C**. T^m/+^ and T^m/LOH^ organoids proliferate at the same rate as measured by CellTiter-Glo luminescent cell viability assay. **D**. T^m/LOH^ organoids are more aggressive *in* vivo than T^m/+^ organoids. C57Bl/6 mice were orthotopically transplanted with the indicated organoid lines and euthanized after 39 days, at which point the mice transplanted with T6^m/LOH^ organoids had reached humane endpoint. The *Trp53* status of the organoid lines transplanted was assessed by PCR-based genotyping. Table of primary tumor occurrences. Hematoxylin and eosin (H&E) staining of tumors or pancreas (if no tumor was observed). Scale bars: 100 μm. **E**. Immunohistochemical staining for p53, p19^Arf^ and γH2AX of tumors formed following orthotopic implantation of T19^m/+^ and T19^m/LOH^ paired organoids.

Once stable cultures were established, T^m/+^ organoids were diploid, while T6^m/LOH^ and T23^m/LOH^ organoids showed a slight increase in DNA content by Propidium Iodide (PI) staining (Fig. 3B) (17). T^m/+^ and T^m/LOH^ organoids proliferated at the same rate *in vitro*, but had different tumorigenic potential following orthotopic implantation in syngeneic mice: the latter forming tumors significantly faster (Fig. 3C, D). Mice transplanted with T^m/+^ organoids developed either no solid tumor or very small tumors when pancreata were collected at the humane endpoint of mice transplanted with the matched T^m/LOH^ organoids. The small tumors derived from mT^m/+^ organoids were moderately or poorly differentiated and showed accumulation of mutant p53, p19^Arf^ and γH2AX, as the tumors derived from mT^m/LOH^ organoids (Fig. 3D, E). Thus, loss of the remaining wild-type *Trp53* allele facilitated tumor growth and the acquisition of invasive features.

### Expression profiling of pre-invasive and invasive tumor organoids identifies transcriptional changes associated with pancreatic cancer progression

To identify the dysregulated pathways following *Trp53* LOH, we performed RNA-seq on 6 pairs of T^m/+^ and T^m/LOH^ organoids. When subjected to PCA, T^m/+^ and T^m/LOH^ organoids separated for the 2 principal components based on their LOH status and paired organoids did not cluster together (Fig. 4A). Detection of the p53 mutation in the RNA-seq data demonstrated that the isolation of *Trp53* LOH cells using Nutlin-3a selection was overall efficient: only 3 lines (T3^m/LOH^, T19^m/LOH^ and T33^m/LOH^) retained few wild-type *Trp53* sequences and were therefore removed from subsequent analyses (Fig. 4B). Comparison of T^m/LOH^ and T^m/+^ organoids identified 1514 significantly differentially expressed genes: 996 up-regulated and 518 down-regulated genes in T^m/LOH^ relative to T^m/+^ (Fig. 4C and Table S2). GSEA revealed, as expected, repression of the p53 signaling pathway in T^m/LOH^ organoids (Fig. 4D). Among the up-regulated genes, we identified several genes that were previously associated with pancreatic cancer progression, such as *Prdm1/*Blimp1, *Foxa1, Gata5, Twist1, Glul* and *Soat1* (Fig. 4E) (11,18,30-32). Thus, these organoid pairs can provide a new platform to identify molecular alterations associated with PDA progression.

**Fig. 4.**
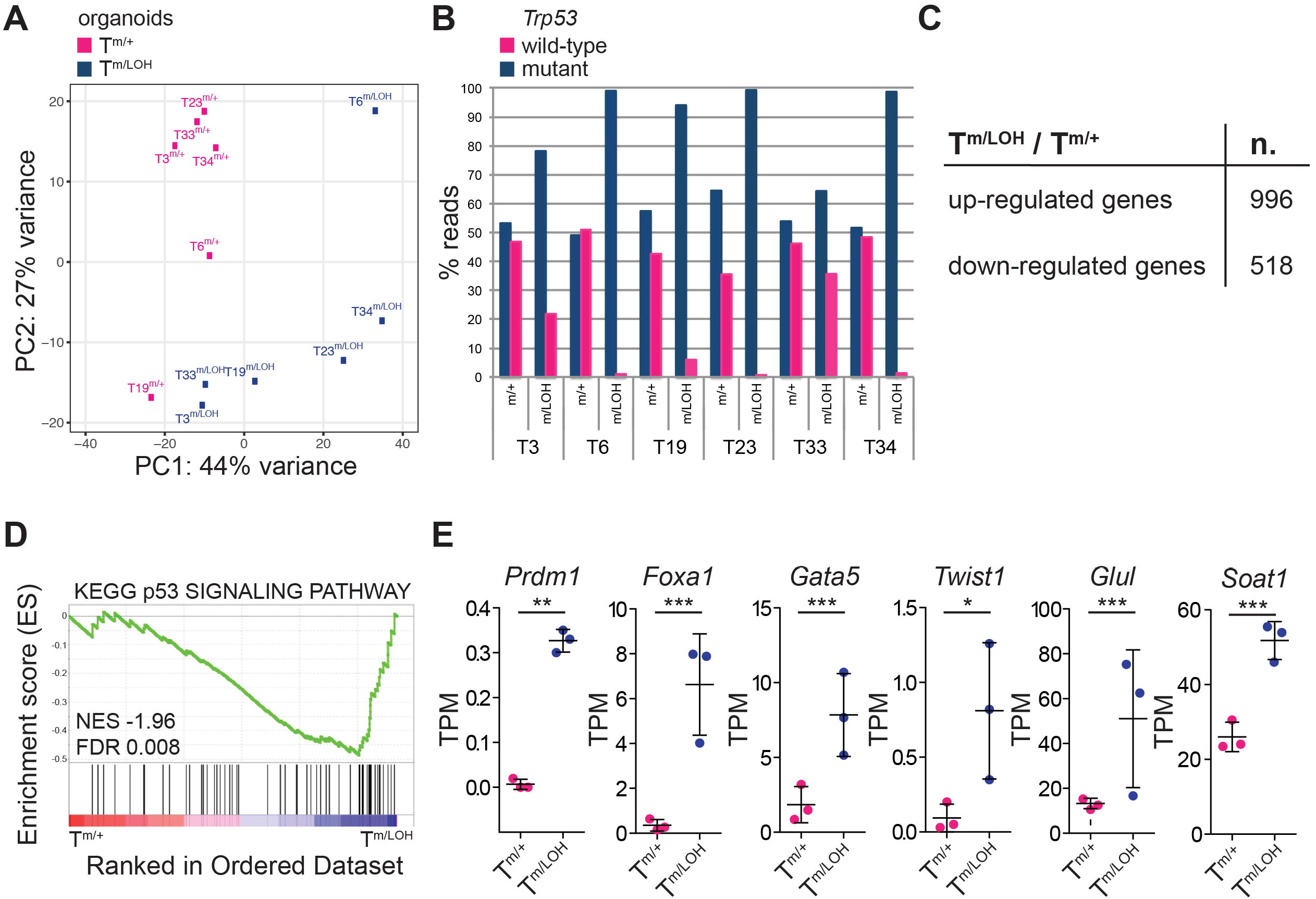
Expression profiling of the tumor organoid pairs provides insights into PDA progression mechanisms. **A**. Principal component analysis of RNA-seq data of T organoid pairs (n=6). Samples are colored based on their group: T^m/+^ in pink and T^m/LOH^ organoids in blue. **B**. Percentage of wild-type and mutant *Trp53* reads as determined by RNA-seq in T^m/+^ (pink bars) and T^m/LOH^ organoids (blue bars). **C**. Table of differentially expressed genes between T^m/LOH^ and T^m/+^ organoids (up-regulated genes: q-value < 0.05, log2 of fold change > 0; down-regulated genes: q-value < 0.05, log2 of fold change < 0). **D**. GSEA results showing that the p53 signaling pathway is repressed in T^m/LOH^ compared to T^m/+^ organoids. NES, normalized enrichment score; FDR, false discovery rate. **E**. *Prdm1, Foxa1, Gata5, Twist1, Glul* and *Soat1* expression in T^m/+^ (pink dots) and T^m/LOH^ organoids (blue dots) as determined by RNA-seq. q-value as calculated by DESeq2 is indicated.

### Fgfr2, Mmp7 and Lcn2 are markers of precursor lesions for pancreatic cancer *in vivo*

Other groups previously reported that PanINs represented the cellular event during pancreatic carcinogenesis that preceded *Trp53* LOH and the acquisition of invasive features (7,33). We sought to identify genes that could be used as biomarkers to detect precursor lesions *in vivo*. Our RNA-seq analysis revealed that *Fgfr2, Mmp7 and Lcn2* were highly expressed in T^m/+^ organoids and almost undetectable in T^m/LOH^ organoids (Fig. 5A, D, G). Next, to evaluate how the expression of these genes was associated with the accumulation of p53 or p19^Arf^ used as invasive markers, we examined their levels by IHC in KPC tumors. We observed that cells co-expressing YFP and Fgfr2 did not show p53 accumulation in KPCY tumor sections (Fig. 5B). In addition, most Fgfr2-positive cells did not present increased levels of p19^Arf^ (Fig. 5C). These cells constituted a small fraction of the tumor mass and represented a histological spectrum of early and advanced PanINs and acinar-to-ductal reprogramming. Our results suggested that Fgfr2 was expressed in precursor lesions of PDA and suppressed in invasive cancer cells exhibiting p53 and p19^Arf^ accumulation.

**Fig. 5.**
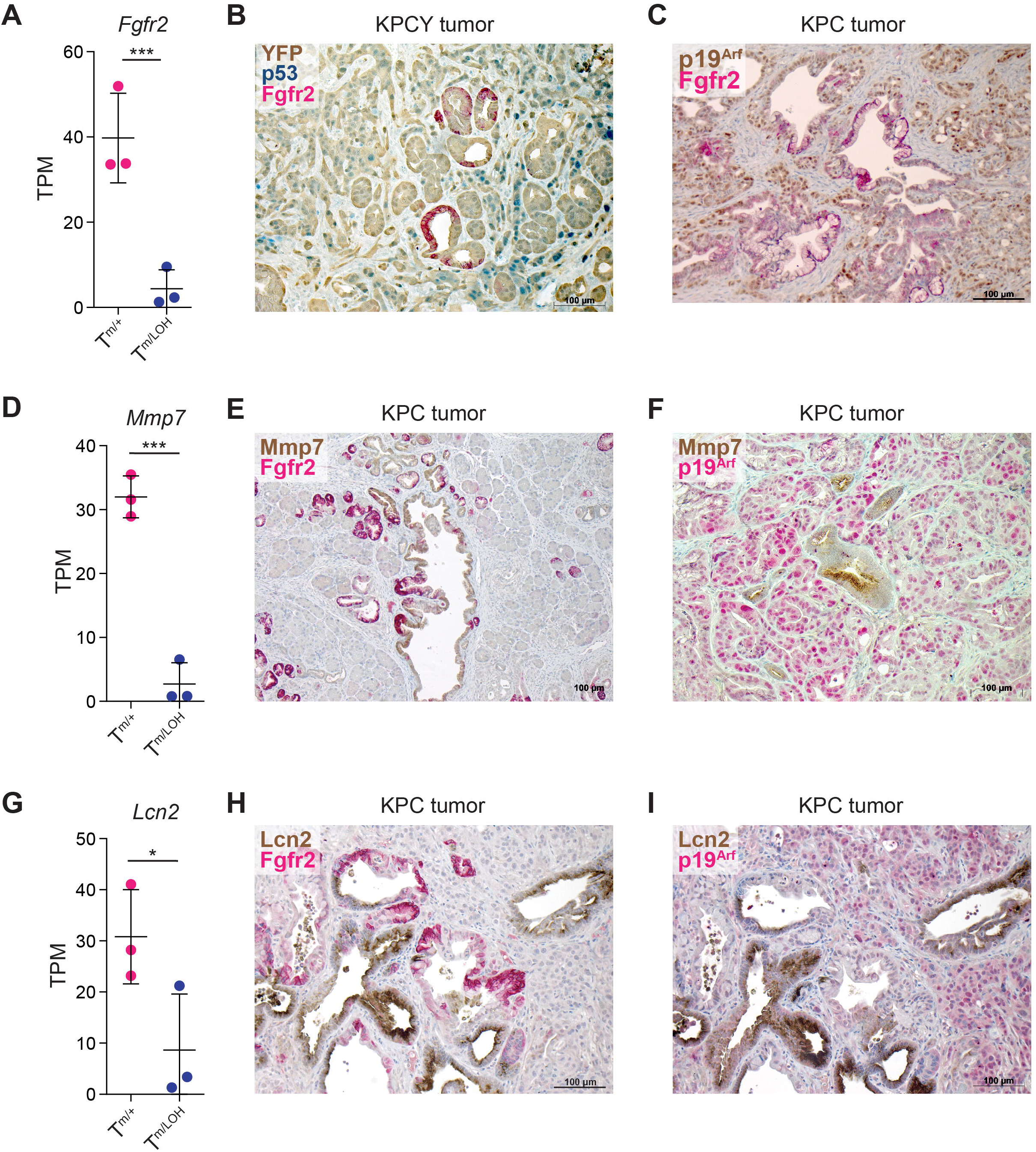
Fgfr2, Mmp7 and Lcn2 are markers of pre-invasive tumor cells *in vivo*. **A, D, G**. *Fgfr2* (**A**), *Mmp7* (**D**) and *Lcn2* (**G**) expression in T^m/+^ (pink dots) and T^m/LOH^ organoids (blue dots) as determined by RNA-seq. q-value as calculated by DESeq2 is indicated. **B**. Immunohistochemical staining for Fgfr2 (red staining), YFP (brown staining) and p53 (blue staining) in a KPCY tumor section. **C**, Immunohistochemical staining for p19^Arf^ (brown staining) and Fgfr2 (red staining) in a KPC tumor section. **E, H**. Immunohistochemical staining for Fgfr2 (red staining) and Mmp7 (**E**) and Lcn2 (**H**) (brown staining) in KPC tumor sections. **F, I**. Immunohistochemical staining for p19^Arf^ (red staining) and Mmp7 (**F**) and Lcn2 (**I**) (brown staining) in KPC tumor sections.

Similarly, *Mmp7* and *Lcn2* were present in PanINs and metaplastic ductal lesions that did also express Fgfr2 and did not show elevated levels of p19^Arf^ (Fig. 5E-F, H-I). It is worth noting that Fgfr2, Mmp7 and Lcn2 expression pattern in precursor lesions was heterogeneous and may have reflected the differentiation state of the pre-invasive cells, which was consistent with data from others that Mmp7 played a role in the initiation and maintenance of metaplastic lesions and Lcn2 in the promotion of PDA progression (34,35). In summary, our results suggested that Fgfr2, Mmp7 and Lcn2 together could serve as the markers to identify PDA precursor cells that retained a functional p53 and constituted the precursor cells for PDA.

### Wild-type p53 favors a premalignant differentiation

Next, we sought to further evaluate whether Fgfr2 was a marker for PDA precursor cells in KPC tumors. Fgfr2-positive PanIN lesions were constituted by columnar cells, which were often organized as a single layer and presented regular nuclei aligned towards the basement membrane (Fig. 6A, B). As neoplastic cells progressed in the disease, they lost polarity, became more disorganized and stopped expressing Fgfr2. Of note, malignant de-differentiation coincided with the acquisition of positive staining for p53 and p19^Arf^, which we have shown by Western blotting to be accumulated in T^m/LOH^ organoids (see Fig. 3A). Additionally, *Trp53* LOH cells presented bigger nuclei (Fig 6A), which were indicative of higher DNA content likely due to aneuploidy, a known consequence of p53 inactivation (36). We observed positive staining for the DNA damage marker γH2AX in Fgfr2-negative and *Trp53* LOH cells (Fig. 6A, C).

**Fig. 6.**
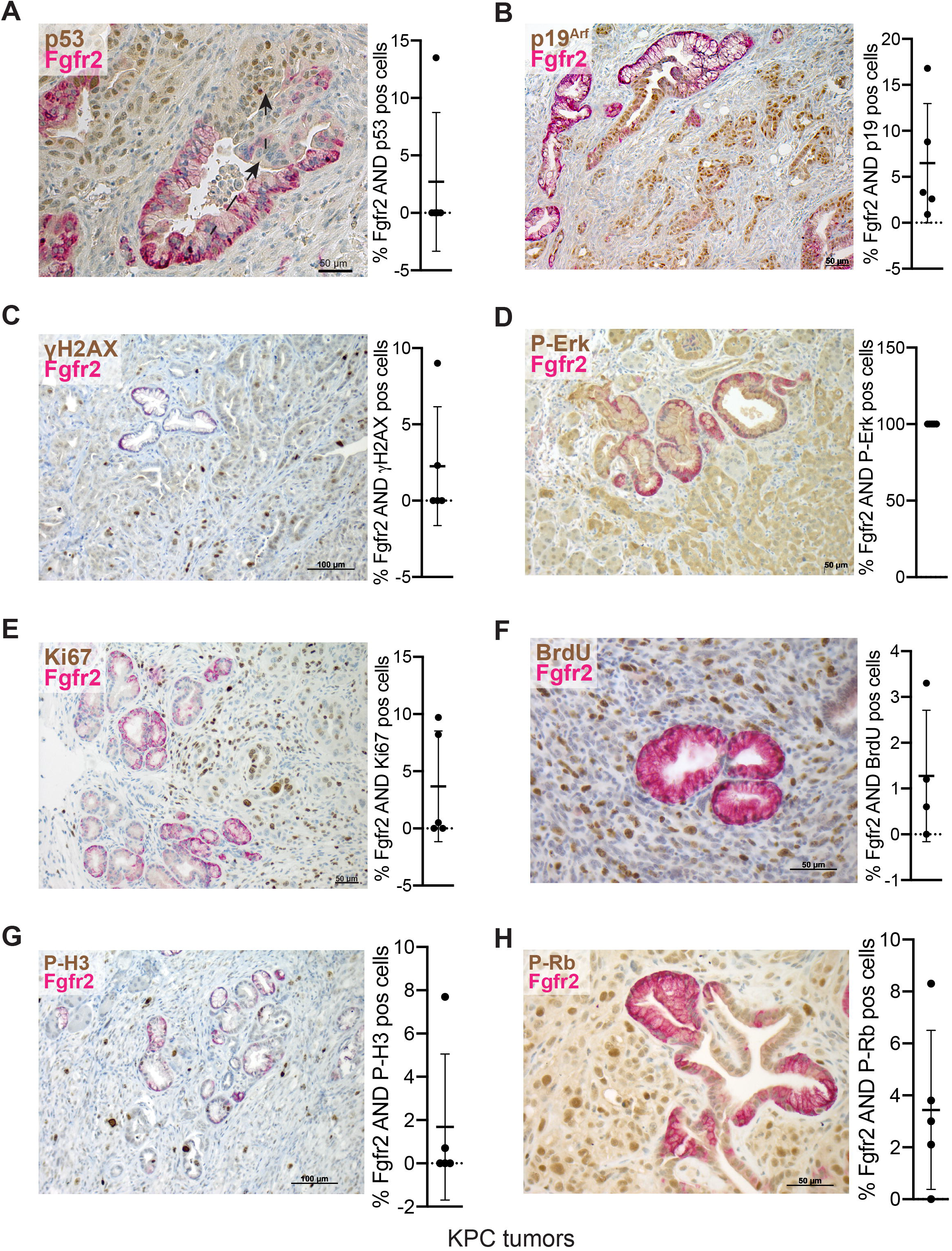
p53 inactivation drives disease progression and the acquisition of invasive features. **A-H**. Left, immunohistochemical staining for Fgfr2 (red staining) and p53 (**A**), p19^Arf^ (**B**), γH2AX (**C**), phospho-p44/42 MAPK (Thr202/Tyr204) (**D**), Ki67 (**E**), BrdU (**F**), phospho-H3 (Ser10) (**G**) and phospho-Rb (Ser807/811) (**H**) (brown staining) in KPC tumor sections. Right, average percentage of double positive cells ± SD. Each dot indicates a tumor section (n=5). The arrow indicates the histological progression of a well-differentiated PanIN lesion to poorly differentiated adenocarcinoma. 10 μl/g of BrdU labeling reagent (Invitrogen) was administered to tumor-bearing mice 24 hr before sample collection.

Of note, cancer cells with and without Fgfr2 expression showed phosphorylation of Erk, indicating that the Kras^G12D^ signaling pathway was active in all neoplastic cells (Fig. 6D). Consistent with findings from other groups that wild-type p53 induced growth arrest in Kras^G12D^-expressing pancreatic cancer cells (13,37,38), most Fgfr2-positive cells did not show markers of proliferation, such as Ki67, phosphorylated histone 3 and phosphorylated Rb, or incorporation of 5*-*bromo-2′*-*deoxy*-*uridine (BrdU) (Fig. 6E-H). Our results supported that wild-type p53 enforced a premalignant differentiation, consistent with a previous report (39). Overall, loss of p53 tumor suppression function was required for cell cycle progression, genomic instability and the acquisition of invasive features, and followed oncogenic Kras-induced metaplastic dysplasia.

## DISCUSSION

In this study, we identified tumor-adjacent pre-invasive cancer cells that expressed Kras^G12D^, p53^R172H^ and one allele of wild-type *Trp53* in the KPC mouse model (Fig. 7). We showed that these cells represented a histological spectrum of early and advanced PanINs and metaplastic lesions, rarely proliferated *in vivo* and did not present markers of advanced disease, such as accumulation of mutant p53, p19^Arf^ and γH2AX. Precursor lesions were constituted by columnar cells, typically organized in a single cell layer. p53 enforced transcriptional programs that maintained the pre-malignant differentiation. Following inactivation of p53, the lesions progressed histologically, underwent malignant de-differentiation and lost polarity, culminating in fully invasive disease.

**Fig. 7.**
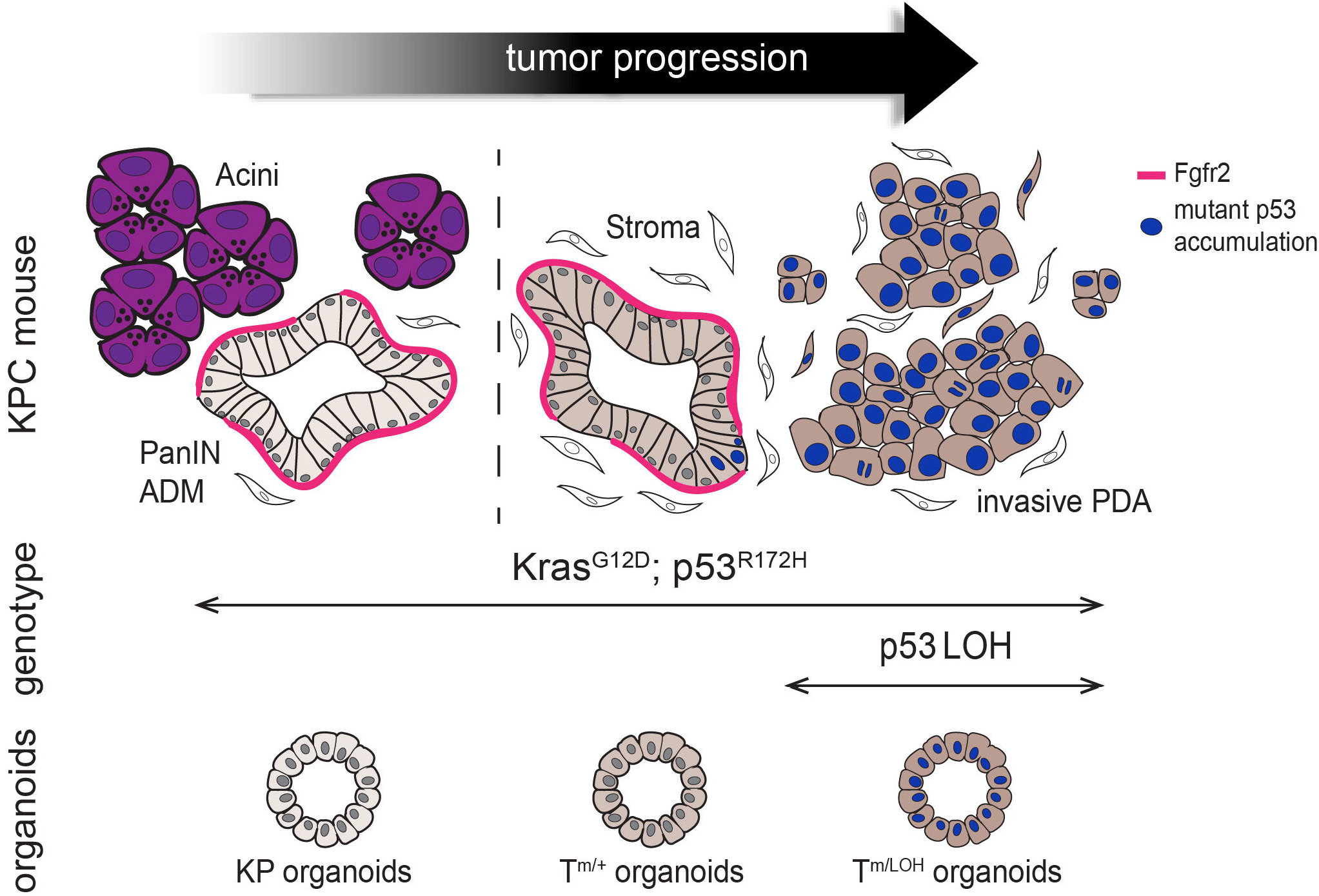
Schematic representation of tumor progression in the KPC mouse model. Activating mutation of Kras in pancreatic acinar or ductal cells initiates precursor lesions, termed PanIN, characterized by progressive cellular and architectural atypia. These pre-invasive cancer cells retain the wild-type *Trp53* allele and express Fgfr2. We have derived KP organoids from low-grade PanINs and T^m/+^ organoids from high-grade PanINs. Following inactivation of p53, pre-invasive cancer cells undergo malignant de-differentiation, start proliferating and acquire markers of advanced disease, such as mutant p53 accumulation, and invasive features.

Using organoid technology, we were able to isolate, expand and characterize matched cultures of pre-invasive primary epithelial tumor cells that retained the wild-type *Trp53* allele (T^m/+^) and invasive tumor cells that had lost the wild-type *Trp53* allele (T^m/LOH^). We showed that pre-invasive tumor cells maintained a diploid genome and grew faster in organoid culture conditions than aneuploid cells. These cells did not present markers of proliferation *in vivo* and had to lose p53 to replicate. *In vitro* they proliferated, possibly induced by the multiple mitogenic factors present in the culture medium. Following orthotopic implantation in syngeneic mice, T^m/+^ cells inactivated the remaining wild-type *Trp53* and formed moderately or poorly differentiated tumors that histologically resembled those formed by invasive cells (T^m/LOH^). Expression profiling of the organoids led to the identification of markers of the pre-invasive cancer cells *in vivo* and mechanisms of disease aggressiveness. In particular, Fgfr2 proved to be a reliable marker of PDA precursor cells in KPC tumors. *Fgfr2b* encodes for a receptor tyrosine kinase whose expression is largely restricted to epithelial cells (40). In KPC tumors, pre-invasive cancer cells manifested epithelial nature and expressed *Fgfr2* (10). Following inactivation of p53, pre-invasive cancer cells lost their epithelial features and underwent malignant de-differentiation (39). Thus, we believe that Fgfr2 marked the epithelial differentiation of pre-invasive cancer cells in KPC tumors.

In summary, this study describes the transition from pre-invasive to invasive PDA at the transcriptional and molecular level and highlights the strengths and the limitations of the organoid system in modeling the disease.

## ACKNOWLEDGEMENTS

We thank Dr. Olaf Klingbeil and Dr. Amber Habowski for critical reading of the manuscript.

This work was performed with assistance from the Cold Spring Harbor Laboratory shared resources, which are supported by the National Institutes of Health (Cancer Center Support Grant 5P30CA045508: Bioinformatics, DNA Sequencing, Flow Cytometry, Animal, and Animal and Tissue Imaging Shared Resources). This work was performed with assistance from the US National Institutes of Health Grant S10OD028632-01.

The authors are supported by National Institutes of Health (Cancer Center Support Grant 5P30CA045508) and the Lustgarten Foundation, where D.A. Tuveson is a distinguished scholar and Director of the Lustgarten Foundation–designated Laboratory of Pancreatic Cancer Research. D.A. Tuveson is also supported by the Thompson Foundation, the Cold Spring Harbor Laboratory and Northwell Health Affiliation, the Northwell Health Tissue Donation Program, the Cold Spring Harbor Laboratory Association, and the National Institutes of Health (5P30CA45508, U01CA210240, R01CA229699, U01CA224013, 1R01CA188134, and 1R01CA190092). This work was also supported by a gift from the Simons Foundation (552716 to D.A. Tuveson). C.

Tonelli was a fellow of the American-Italian Cancer Foundation. Y. Park is supported by the National Cancer Institute (R50CA211506).

## FIGURE LEGENDS

**Suppl. Fig. 1.**
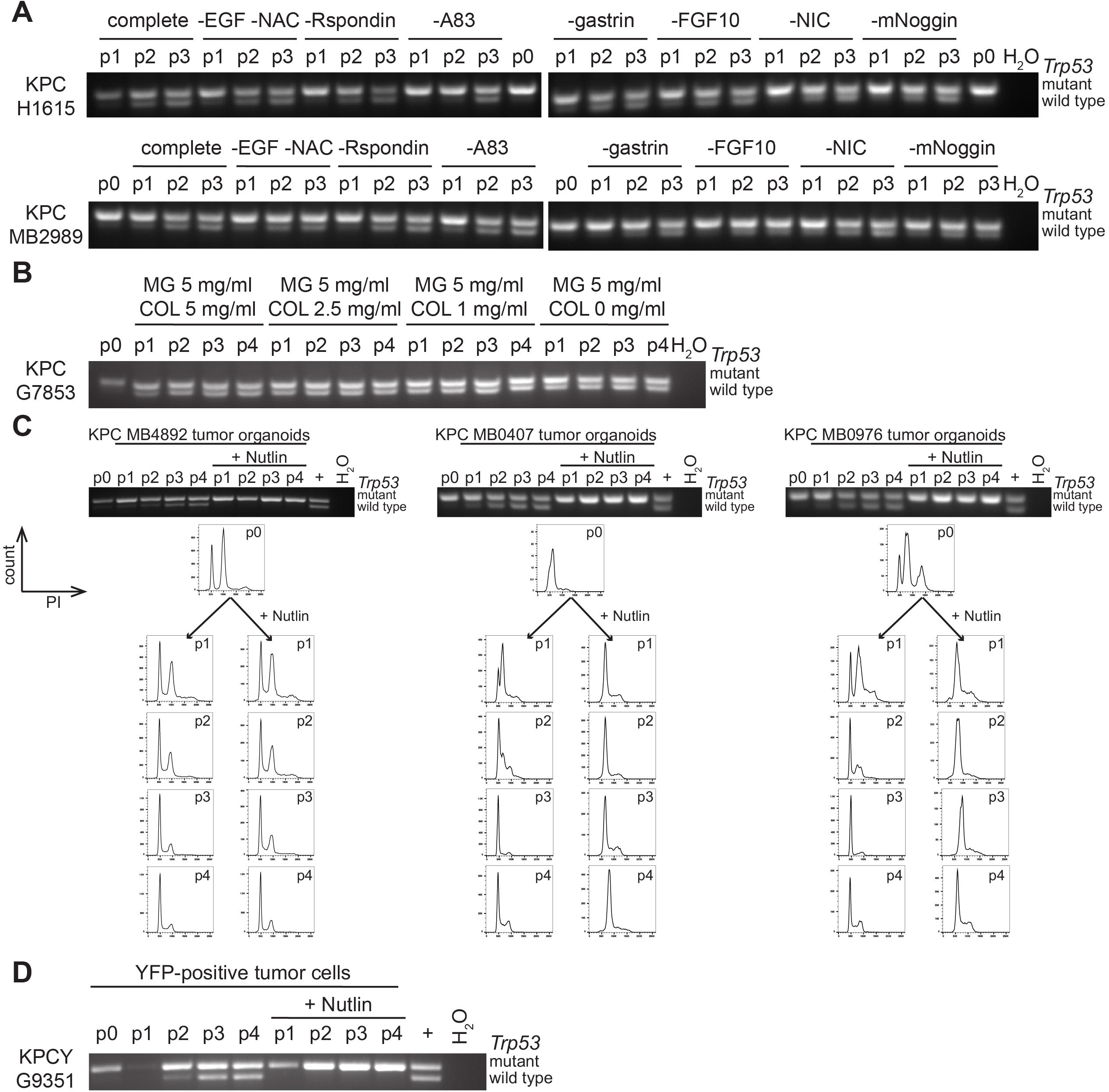
**A**. PCR-based genotyping for assessing the *Trp53* status in tumor cells from KPC mice after isolation (p0) and upon passaging as organoids in medium depleted of different components (p1 to p3). **B**. PCR-based genotyping for assessing the *Trp53* status in tumor cells from a KPC mouse after isolation (p0) and upon passaging as organoids in matrigel supplemented with different amounts of collagen (p1 to p4). **C**. PCR-based genotyping for assessing the *Trp53* status in tumor cells from KPC mice after isolation (p0) and upon passaging as organoids in medium with or without Nutlin-3a (p1 to p4). The DNA profile of the tumor cells was evaluated at every passage by PI staining. **D**. PCR-based genotyping for assessing the *Trp53* status in FACS-sorted tumor cells from a KPCY mouse after isolation (p0) and upon passaging as organoids in medium with or without Nutlin-3a (p1 to p4).

**Suppl. Fig. 2.**
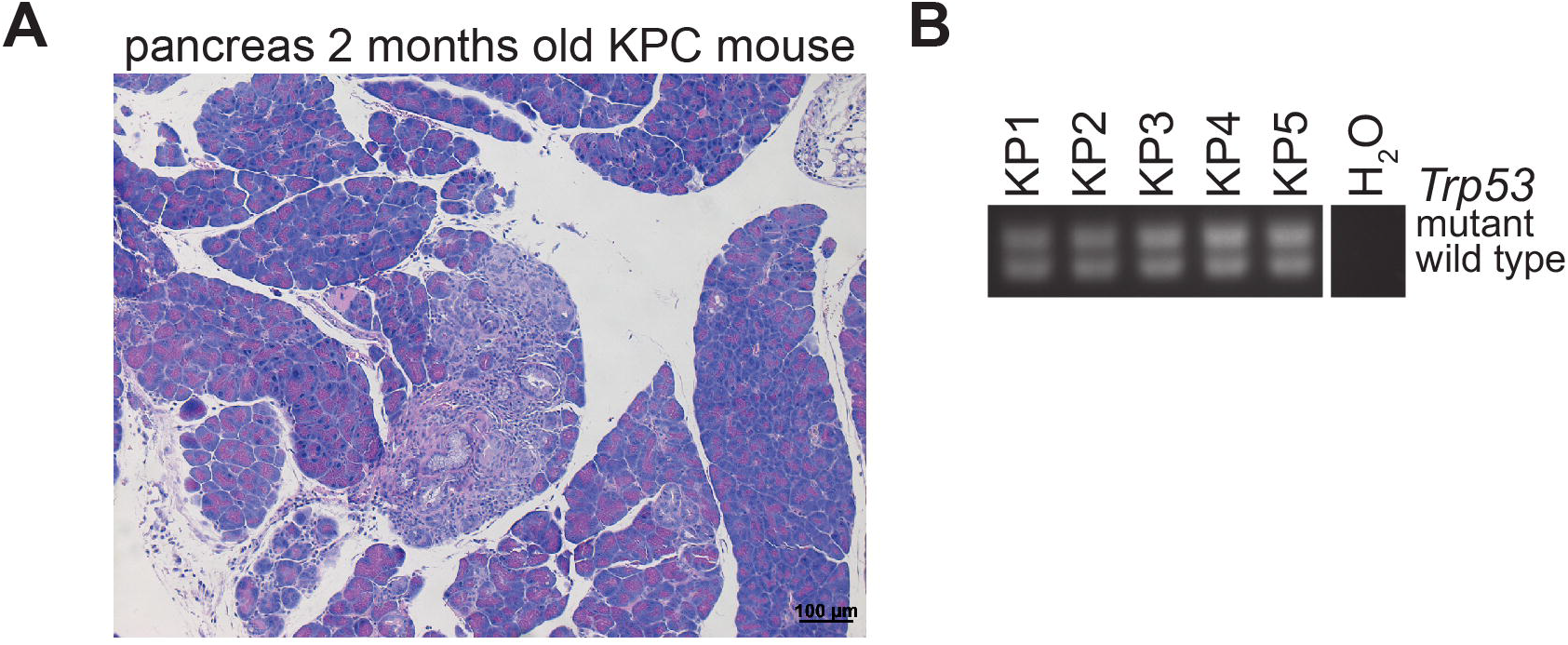
**A**. Representative H&E staining of the pancreas of a 2-month-old KPC mouse. **B**. PCR-based genotyping for assessing the *Trp53* status in KP organoids.

## MATHERIALS & METHODS

### Animals

*Kras*^*LSLG12D*^, *Trp53*^*LSLR172H*^, *Pdx1-Cre* and *Rosa26*^*LSLYFP*^ strains in C57Bl/6 background were interbred to obtain *Kras*^*LSLG12D/+*^; *Trp53*^*LSLR172H/+*^; *Pdx1-Cre* (KPC) and *Kras*^*LSLG12D/+*^; *Trp53*^*LSLR172H/+*^; *Pdx1-Cre; Rosa26*^*LSLYFP*^ (KPCY) mice (7,10,12). C57Bl/6 were bred in house. All animal experiments were conducted in accordance with procedures approved by the IACUC at Cold Spring Harbor Laboratory (CSHL).

### Tumor cells isolation

Primary tumor tissues were carefully dissected, avoiding adjacent normal pancreas or other tissue contamination. Tumors were minced and digested for 1 hr at 37°C in digestion buffer (DMEM, 5% FBS, penicillin, streptomycin, 2.5 mg/ml Collagenase D (Sigma), 0.5 mg/ml Liberase DL (Sigma), 0.2 mg/ml DNase I (Sigma)) while shaking. Single-cell suspensions were obtained by filtering through 100 μm nylon cell strainers and subsequent hypotonic lysis of red blood cells using ACK lysis buffer (Gibco). KPC tumor cells were enriched using mouse Tumor Cell Isolation Kit (Miltenyi Biotech) by magnetic cell sorting (MACS), according to manufacturer’s instructions. To isolate KPCY tumor cells, cells were stained with 1 μg/ml DAPI and DAPI-negative YFP-positive cells were sorted with the FACSAria cell sorter (BD). Tumor cells were prepared for subsequent experiments or seeded in growth-factor-reduced (GFR) Matrigel (BD).

### Murine Pancreatic Ductal Organoid Culture

Tumor organoids were established as described in (17). T^m/+^ and T^m/LOH^ organoid pairs were derived from passage 1 organoids (17,18). Cells were seeded in GFR Matrigel (BD). Once Matrigel was solidified, pancreatic organoid medium was added. Pancreatic organoid medium contains AdDMEM/F12, 10 mM HEPES (Invitrogen), Glutamax 1X (Invitrogen), penicillin/streptomycin 1x (Invitrogen), 500 nM A83-01 (Tocris), 50 ng/ml mEGF, 100 ng/ml mNoggin (Peprotech), 100 ng/ml hFGF10 (Peprotech), 0.01 mM hGastrin I (Sigma), 1.25 mM N-acetylcysteine, 10 mM Nicotinamide (Sigma), B27 supplement 1X (Invitrogen), R-spondin conditioned medium (10% final) and 10 μM Nutlin-3a (Sigma) when indicated. T^m/LOH^ organoids were passaged 3 times in organoid medium with 10 μM Nutlin-3a (Sigma).

For dissociating organoids into single cell suspensions, organoids were incubated in TrypLE Express Enzyme (ThermoFisher) for 15 min while shaking. To generate KPC-2D cell lines from tumor organoid cultures, organoids were dissociated into single cells as described above, resuspended with DMEM supplemented with 5% FBS, penicillin and streptomycin and plated on tissue culture plates. Bright Field pictures of organoids were taken with a Nikon eclipse TE2000-S microscope.

### PCR-based genotyping of *p53* 1loxP

Organoids were harvested from two wells of 24 well plate and centrifuged at 4000rpm for 5 min at 4C. Genomic DNA from freshly isolated tumor cells or organoids was extracted with DNEasy Blood & Tissue Kit (Qiagen) following the protocol for cultured cells. Each PCR reaction for p53 1loxP genotyping was performed in a 20 μl mixture containing 1x AmpliTaq Gold 360 master mix (ThermoFisher), 0.5 μM each primer and 40 ng template DNA. The following primers were used for genotyping: *For* AGCCTGCCTAGCTTCCTCAGG, *Rev* CTTGGAGACATAGCCACACTG. The PCR cycling conditions were 95°C for 5 minutes, followed by 40 cycles at 95°C for 30 seconds, 56°C for 30 seconds, and 72°C for 30 seconds, with a final extension step at 72°C for 5 minutes. PCR products were separated on a 2% agarose gel in 1x TAE buffer. Gel imaging was performed with a Syngene UV transilluminator.

### DNA content analysis by Propidium Iodide staining

Organoids were harvested from two wells of 24 well plate and dissociated into single cells by incubating them in TrypLE Express Enzyme (ThermoFisher) for 15 min while shaking. To analyze DNA content profile, 1×10^5^ freshly isolated tumor cells or organoid cell suspensions were resuspended in 1 ml of PBS and fixed by adding 2 ml of ice-cold absolute ethanol and kept at 4°C for at least 30 minutes. Cells were washed once with 1 ml of 1% BSA in PBS and stained overnight with 50 μg/ml Propidium Iodide and 250 μg/ml RNaseA at 4°C. All fluorescence-activated cell sorting (FACS) data were acquired using a LSRFortessa cell analyzer (BD) and analyzed with FlowJo software (TreeStar).

### *In vitro* proliferation assay on mouse organoids

Organoids were dissociated into single cells by incubating them in TrypLE Express Enzyme (ThermoFisher) for 15 min while shaking. Cells were counted and diluted to 10 cells/μl in a mixture of pancreatic organoid medium (90% final concentration) and GFR Matrigel (BD, 10% final concentration). 150 μl per well of this mixture (1500 cells per well) was plated in 96-well white plates (Nunc), whose wells had been previously coated with poly(2-hydroxyethyl methacrylate) (Sigma) to prevent cell adhesion to the bottom of the wells. Cell viability was measured every 24 hr, starting one day after plating, using the CellTiter-Glo assay (Promega) and SpectraMax I3 microplate reader (Molecular Devices). Four replicate wells per time point were measured.

### Western blotting

For western blotting experiments with PDA organoids, organoids were harvested from six wells of 24 well plate. Cells were lysed with RIPA Buffer (300 mM NaCl, 5 mM EDTA, 20mM HEPES, 10% glycerol, 1% Triton X-100) supplemented with protease inhibitors (cOmplete, Mini, EDTA-free Protease Inhibitor Cocktail, Roche) and phosphatase inhibitors (PhosSTOP, Roche). Cleared lysates were electrophoresed and immunoblotted with the indicated primary antibodies: HSP90 (EMD Millipore, 07-2174), cyclin G1 (ThermoFisher, PA5-36050), p53 (Santa Cruz, FL393), p19^Arf^ (Abcam, ab26696), phospho-Histone H2A.X (Ser139) (CST, 9718 clone 20E3). After incubation of the membranes with appropriate HRP-conjugated secondary antibodies (Jackson ImmunoResearch Laboratories), imaging was performed using an enhanced chemiluminescence (ECL) detection kit (Sigma) and autoradiography films (VWR, 165 1454).

### *In vivo* transplantation assay

Organoids were cultured as described above and quickly harvested on ice in AdDMEM/F12 medium supplemented with HEPES 1x (Invitrogen), Glutamax 1x (Invitrogen), and penicillin/streptomycin (Invitrogen). Organoids were dissociated to single cells with TrypLE Express Enzyme (ThermoFisher). Cells were resuspended in 50 μl of GFR Matrigel (BD) diluted 1:1 with cold PBS. Mice were anesthetized using Isoflurane and subcutaneous administration of Ketoprofen (5 mg/kg). The surgery site was disinfected with Iodine solution and 70% ethanol. An incision was made in the upper left quadrant of the abdomen. 10^5^ cells were injected in the tail region of the pancreas of wild-type C57Bl/6 mice. The incision at the peritoneal cavity was sutured with Coated Vicryl suture (Ethicon) and the skin was closed with wound clips (Reflex7, CellPoint Scientific Inc). Mice were euthanized 39 days after surgery. Tumors were dissected out and processed for histology.

### Histology

All tissues were fixed with 10% neutral buffered formalin overnight. Tissues were then processed with standard tissue processing protocol, embedded in paraffin and 6 μm sections were cut and mounted on slides. Formalin fixed paraffin-embedded tissue sections were stained with hematoxylin and eosin or used for immunohistochemical staining.

### Immunohistochemistry

FFPE sections were deparaffinized and rehydrated. For antigen retrieval, slides were incubated with 10 mM Citrate buffer (pH 6.0) in a pressure cooker for 6 minutes. To perform immunohistochemistry (IHC), endogenous peroxidase activity was quenched in 3% H_2_O_2_ for 20 min. Tissues were blocked in 2.5% Normal Horse or Goat Serum blocking solution (Vector Laboratories) for IHC and subjected to staining with the following antibodies overnight at 4°C: p53 (Leica, P53-CM5P-L 1:100), GFP/YFP (Abcam, ab6673 1:100), p19^Arf^ (Abcam, ab26696 1:1000), phospho-Histone H2A.X (Ser139) (CST, 9718 clone 20E3 1:1000), Fgfr2 (Abcam, ab58201 1:1000), phospho-p44/42 MAPK (Erk1/2) (Thr202/Tyr204) (CST, 4370 clone D13.14.4E 1:250), Ki67 (ThermoFisher, RM-9106-S clone SP6 1:250), BrdU (Abcam, ab152095 1:500), phospho-Histone H3 (Ser10) (CST, 9701 1:200), phospho-Rb (Ser780) (CST, 9307 1:100), Mmp7 (CST, 3801 clone D4H5 1:100), Lipocalin-2 (Abcam, ab63929 1:100). ImmPRESS Alkaline Phosphatase (AP) anti-Rat IgG polymer, ImmPRESS AP anti-Mouse IgG polymer, ImmPRESS AP anti-Rabbit IgG polymer and ImmPRESS Horseradish Peroxidase (HRP) anti-Rabbit IgG polymer, ImmPRESS HRP anti-Rat IgG polymer (Vector Laboratories) were used as secondary antibodies for IHC. ImmPACT DAB peroxidase substrate, Vector Blue Alkaline Phosphatase substrate and Vector Red Alkaline Phosphatase substrate (Vector Laboratories) were used as substrates. Hematoxylin (Vector Laboratories) was used as counterstain. Cover slides were mounted with Cytoseal 60. Brightfield images were obtained using a Zeiss Axio Imager.A2 microscope.

### RNA-sequencing of Murine Organoids

For each organoid line, 4-6 wells of organoids from a 24 well plate were harvested in 1 ml of TRIzol reagent (ThermoFisher) and flash-frozen. RNA was extracted using the TRIzol Plus RNA Purification Kit (ThermoFisher) per manufacturer’s instructions. RNA samples were treated on column with PureLink DNase (ThermoFisher). The quality of purified RNA samples was determined using a Bioanalyzer 2100 (Agilent) with an RNA 6000 Nano Kit. RNAs with RNA Integrity Number (RIN) values greater than 8.5 were used to generate sequencing libraries. Libraries were generated from 1 μg of total RNA using a TruSeq Stranded Total RNA Library Prep Human/Mouse/Rat (48 Samples) (Illumina) per manufacturer’s instructions. Libraries were quality checked using a Bioanalyzer 2100 (Agilent) with a High Sensitivity DNA Kit and quantified using PicoGreen (ThermoFisher). Equimolar amounts of libraries were pooled and subjected to paired-end, 75 bp sequencing at the Cold Spring Harbor DNA Sequencing Next Generation Shared Resource using an Illumina NextSeq 500 platform.

### RNA-sequencing Data Analysis

RNA-seq reads quality was first quantified using FastQC (http://www.bioinformatics.babraham.ac.uk/projects/fastqc/). Reads were mapped to transcript annotation GENCODE M16 (41) using Spliced Transcripts Alignment to a Reference (STAR) (42). RSEM (43) was used to extract counts per gene and transcripts per million (TPM).

RNA-seq tracks were generated from aligned reads using deepTools2 (44) and visualized using UCSC Genome Browser (45).

Differential gene expression analysis was performed using Bioconductor package DESeq2 (46). T^m/+^ and T^m/LOH^ organoids were considered paired datasets, while KP and T^m/+^ organoids unpaired datasets. A pre-filtering step was performed to remove genes that have reads in less than two samples. At this step, all genes not classified as protein coding according to BioMart were discarded (47). Only genes with an adjusted p-value less than 0.05 were retained as significantly differentially expressed genes. A principal component analysis (PCA) was performed using the 500 most variable protein coding genes after variance stabilizing transformation (VST) using DESeq2 software. The graphical representation of the two most important components was created using CRAN package ggplot2 (48).

Default parameters were used to perform GSEA Preranked analysis using the GSEA software (49,50). Hallmark and curated gene sets were downloaded from the MSigDB database, which include pathways from the Kyoto Encyclopedia of Genes and Genomes (KEGG) (51,52).

